# Geochemical influences on nonenzymatic oligomerization of prebiotically relevant cyclic nucleotides

**DOI:** 10.1101/872234

**Authors:** Shikha Dagar, Susovan Sarkar, Sudha Rajamani

## Abstract

The spontaneous emergence of RNA on the early Earth continues to remain an enigma in the field of origins of life. Few studies have looked at the nonenzymatic oligomerization of cyclic nucleotides under neutral to alkaline conditions, in fully dehydrated state. Herein, we systematically investigated the oligomerization of cyclic nucleotides under prebiotically relevant conditions, where starting reactants were subjected to repeated dehydration-rehydration (DH-RH) regimes, like they would have been on an early Earth. DH-RH conditions, a recurring geological theme, are driven by naturally occurring processes including diurnal cycles and tidal pool activity. These conditions have been shown to facilitate uphill oligomerization reactions in terrestrial geothermal niches, which are hypothesized to be pertinent sites for the emergence of life. 2′-3′ and 3′-5′ cyclic nucleotides of one purine-based (adenosine) and one pyrimidine-based (cytidine) system were evaluated in this study. Additionally, the effect of amphiphiles was also investigated. Furthermore, to discern the effect of ‘realistic’ conditions on this process, the reactions were also performed using hot spring water samples from an early Earth analogue environment. Our results showed that the oligomerization of cyclic nucleotides under DH-RH conditions resulted in intact informational oligomers. Amphiphiles increased the stability of, both, the starting monomers and the resultant oligomers. In analogue condition reactions, oligomerization of nucleotides and back-hydrolysis of the resultant oligomers was pronounced. Altogether, this study demonstrates how nonenzymatic oligomerization of cyclic purine and pyrimidine nucleotides, under laboratory-simulated and early Earth analogous conditions, could have resulted in RNA oligomers of a putative RNA World.

## 1. Introduction

The “RNA World hypothesis” emphasizes the potential of RNA molecules to be the first biomolecule to have emerged on the prebiotic Earth due to its capability to act as a catalyst, in addition to being able to encode genetic information (Gilbert 1986). The presence of large complex enzymes assisting the process of encoding and processing genetic information would have been implausible on the early Earth. Therefore, the formation and replication of RNA molecules on the early Earth would have been nonenzymatic, and driven by physico-chemical properties of molecules, resulting in short oligonucleotides. Such small oligoribonucleotides have been shown to not only enhance the rate of template-directed replication (Kaiser and Richert 2013; Li et al. 2017; Tam et al. 2017; O’Flaherty et al. 2018), but are also thought to act as cofactor precursors that pervades extant biology (Puthenvedu et al. 2015; Majerfeld et al. 2016; Puthenvedu et al. 2017).

Previous studies that explored the formation and replication of RNA molecules under prebiotic conditions, typically used chemically modified nucleotides such as imidazole-activated nucleotides, as the starting monomers (Lohrmann and Orgel 1973; Blain and Szostak 2014; Puthenvedu et al. 2015; Majerfeld et al. 2016; Puthenvedu et al. 2017). Though a recent study has demonstrated the synthesis of such imidazole-activated nucleotides under prebiotic conditions (Yi et al. 2018), their availability in significant amounts is still uncertain. Furthermore, the extreme susceptibility of these activated nucleotides towards high temperatures and aqueous conditions, make their potential role as substrates for enzyme-free oligomerization highly debatable. On the other hand, the lipid-assisted nonenzymatic oligomerization of nucleoside 5′-monophosphates (NMP) has been demonstrated, under terrestrial acidic geothermal conditions (Rajamani et al. 2008). Nevertheless, systematic characterization of these products showed the formation of RNA-like polymers with abasic sites (Mungi et al. 2015), due to the loss of the informational moiety under the aforesaid conditions. Hence, it is disputable that the resultant moieties which formed under such conditions, would have been capable of functioning as informational oligomers. Conversely, cyclic nucleotides can readily form under prebiotic conditions and are comparatively more stable while being intrinsically active (Tapiero and Nagyvary 1971; Verlander et al. 1973; Verlander and Orgel 1974; Powner et al. 2009; Costanzo et al. 2007; Powner et al. 2010; Saladino et al. 2012; Suárez-Marina et al. 2019). Therefore, they could have potentially served as monomers for enzyme-free oligomerization, to result in RNA. Towards this, Verlander et al. demonstrated the nonenzymatic oligomerization of 2′-3′ cyclic nucleoside monophosphates (cNMPs) under completely dry conditions (Verlander et al. 1973). Subsequent studies from di Mauro’s group showed the formation of RNA oligomers using 3′-5′ cyclic nucleotides under slightly alkaline regimes and in dry conditions (Costanzo et al. 2012a, 2016b). Although insightful, an important aspect pertaining to these studies is the uncertainty of the existence of permanently dry niches on the early Earth. The realistic approach would then be to consider these reactions in niches that have been hypothesized to support the chemical emergence of life. One of these proposed niches is that of terrestrial hydrothermal pools. Temperature and seasonal fluctuations in these terrestrial pools, could lead to dry-wet cycles (Dehydration-Rehydration/DH-RH cycles) (Higgs 2016; Ross and Deamer 2016), a feature very common to the geology of the planet. The dry phase can concentrate the starting monomers, thus facilitating concentration-dependent uphill reactions such as oligomerization. Subsequent wet phase could provide a medium for the diffusion of the growing oligomers, albeit favoring back reactions like hydrolysis also. Previous studies have demonstrated the formation of oligoesters and depsipeptides, under DH-RH conditions by condensation of hydroxy acids alone and in combination with amino acids, respectively (Mamajanov et al. 2014; Mamajanov 2015). As mentioned earlier, lipids have been shown to facilitate the oligomerization of 5′-NMPs under DH-RH conditions, resulting in the formation of RNA-like oligomers (abasic oligomers) (Rajamani et al. 2008; Mungi et al. 2015). Upon dehydration in these scenarios, lipids are thought to get arranged in two-dimensional arrays, concentrating the reacting monomers within the interlamellar spaces, thereby promoting concentration-dependent oligomerization reactions (Damer and Deamer 2015). Upon rehydration, the multilamellar structures can encapsulate the monomers, as well as the growing oligomers, forming protocell-like entities, thus protecting them from hydrolysis. Nonetheless, lipid-assisted nonenzymatic oligomerization of cyclic nucleotides, under aforesaid prebiotic conditions, has not yet been demonstrated. Therefore, in this study, we also aimed to discern the effect of lipids on the oligomerization of cyclic nucleotides, under prebiotically pertinent DH-RH regimes, to discern how their oligomerization would have been facilitated in the terrestrial hydrothermal pools of the early Earth.

Most of the previous studies undertaken to demonstrate the nonenzymatic oligomerization of cyclic nucleotides have used purines as the starting monomer (Verlander et al. 1973; Verlander and Orgel 1974; Pino et al. 2008; Costanzo et al. 2016a). Very few studies have looked at the oligomerization of pyrimidines under dry conditions (Tapiero and Nagyvary 1971; Costanzo et al. 2017). Purines are known for their tendency to stack better, and thus are thought to oligomerize more efficiently than pyrimidines (Giovanna et al. 2009). Stacking increases the proximity of the nucleotides, thus facilitating intermolecular condensation that results in oligomerization. Given this, we wanted to elucidate whether there is any effect from the aforesaid difference (purine vs pyrimidine) on the oligomerization of cyclic nucleotides under DH-RH conditions. Towards this, enzyme-free oligomerization of cyclic nucleotides was studied using cyclic adenosine monophosphate (cAMP), a purine, and cyclic cytosine monophosphate (cCMP), a pyrimidine nucleotide, using both the 2′-3′ and 3′-5′ cyclic isomer versions. Lastly, most of the studies pertaining to discerning the chemical origins of life are constrained to reactions that are carried out under laboratory-controlled conditions. Although insightful, such controlled reactions may not be representative of the complex scenarios that might have been present on the early Earth. Deamer et. al. tried to constrain the range of possible environments/conditions for the origins of life by analyzing prebiotically pertinent prebiotic processes under realistic conditions (Deamer et al. 2006). Previous study in our lab have demonstrated the self-assembly of simple amphiphiles, like fatty acids and their derivatives, in alkaline hot spring water samples of varying ionic content that were collected from three locations in Ladakh, an early Earth analogue site (Joshi et al. 2017). However, to our knowledge, similar analogue environment studies have not been undertaken in the context of nonenzymatic oligomerization of nucleotides to form RNA. Therefore, we also set out to investigate how cyclic nucleotide oligomerization reactions would advent if the reactions were to be carried out in a complex ‘realistic’ context, instead of a controlled laboratory set-up by using the same hot spring sample that were used in the previous study (Joshi et al. 2017).

The present study reports the nonenzymatic oligomerization reactions of cyclic nucleotides under DH-RH regimes in laboratory simulated conditions as well as in an early Earth analogue environment. Under the laboratory simulated conditions, intact oligomers up to trimers (in pyrimidine-based reactions) and tetramers (in purine-based reactions) were observed in some of the oligomerization reactions. Interestingly, in the reactions performed under analogue conditions, the hot-spring water sample used, seemed to enhance the rate of oligomerization, but also led to the destabilization of the oligomers. Additionally, presence of lipid seemed to enhance the stability of starting monomers as well as oligomers formed.

## 2. Results

### 2.1. Oligomerization of cyclic adenosine monophosphate under DH-RH conditions

As alluded to earlier, DH-RH cycles are thought to have been widely prevalent in terrestrial geothermal environments. Therefore, we investigated the enzyme-free oligomerization of cyclic nucleotides under such fluctuating conditions; a niche that has been shown to promote the accumulation of increased oligomerization products over multiple cycles (Rajamani et al. 2008; Mamajanov et al. 2014; Mamajanov 2015; Mungi et al. 2015). A typical reaction involved 5 mM of the corresponding cyclic nucleotides (2′-3′ or 3′-5′ cNMP) at pH 8, which were subjected to thirty DH-RH cycles (with 24 hours per cycle) (Method 4.2.1 and 4.2.3). A dry heating control reaction was also set up under similar conditions, wherein the sample was only subjected to prolonged drying without any rehydration event involved. This was also done to recreate conditions used in previous studies (Verlander et al. 1973; Verlander and Orgel 1974; Costanzo et al. 2012a, 2016a). The reactions were analyzed using High Performance Liquid Chromatography (HPLC) (Method 4.2.5). The product identities were ascertained by co-eluting them on HPLC with known control samples. The control species including cAMP, AMP monomer, dimer, and trimer peaks eluted at 1.8 minutes, 6.1 minutes, 6.3 minutes and 7.1 minutes, respectively. The peak observed with a retention time of 1.5 minutes is adenine, which is mentioned as a breakdown peak in the chromatogram (Figure 1, Panel A). As free adenine does not interact with an anion exchange column like the one used for this analysis, it elutes out of the column in void volume. As shown in the HPLC chromatogram (Figure 1, Panel B), the reaction mixture kept at 90 °C for thirty days under dry heating conditions led to the hydrolysis of cAMP to its linear form (i.e. AMP), no oligomers were observed. Conversely, under DH-RH conditions, a peak, which had similar retention time as that of the control dimer, was observed in the chromatogram corresponding to 2′, 3′ cAMP oligomerization reaction (Figure 1, Panel A). Nonetheless, there was no clear separation observed among the other peaks with a retention time more than that of AMP, in the HPLC chromatogram. This could be, both, due to the relatively low yields of the reaction, and due the presence of different types of product species. For e.g., resultant products could be either cyclic or linear versions, and could also have varied internucleotide linkages, especially in the 2′, 3′ cNMP reactions. However, in the case of 3′, 5′ cAMP, oligomeric species was not detected with HPLC for both the control and the DH-RH reaction sets (Figure 1, Panels C and D). All the reactions were performed at least in replicate to validate the results.

**Figure 1:**
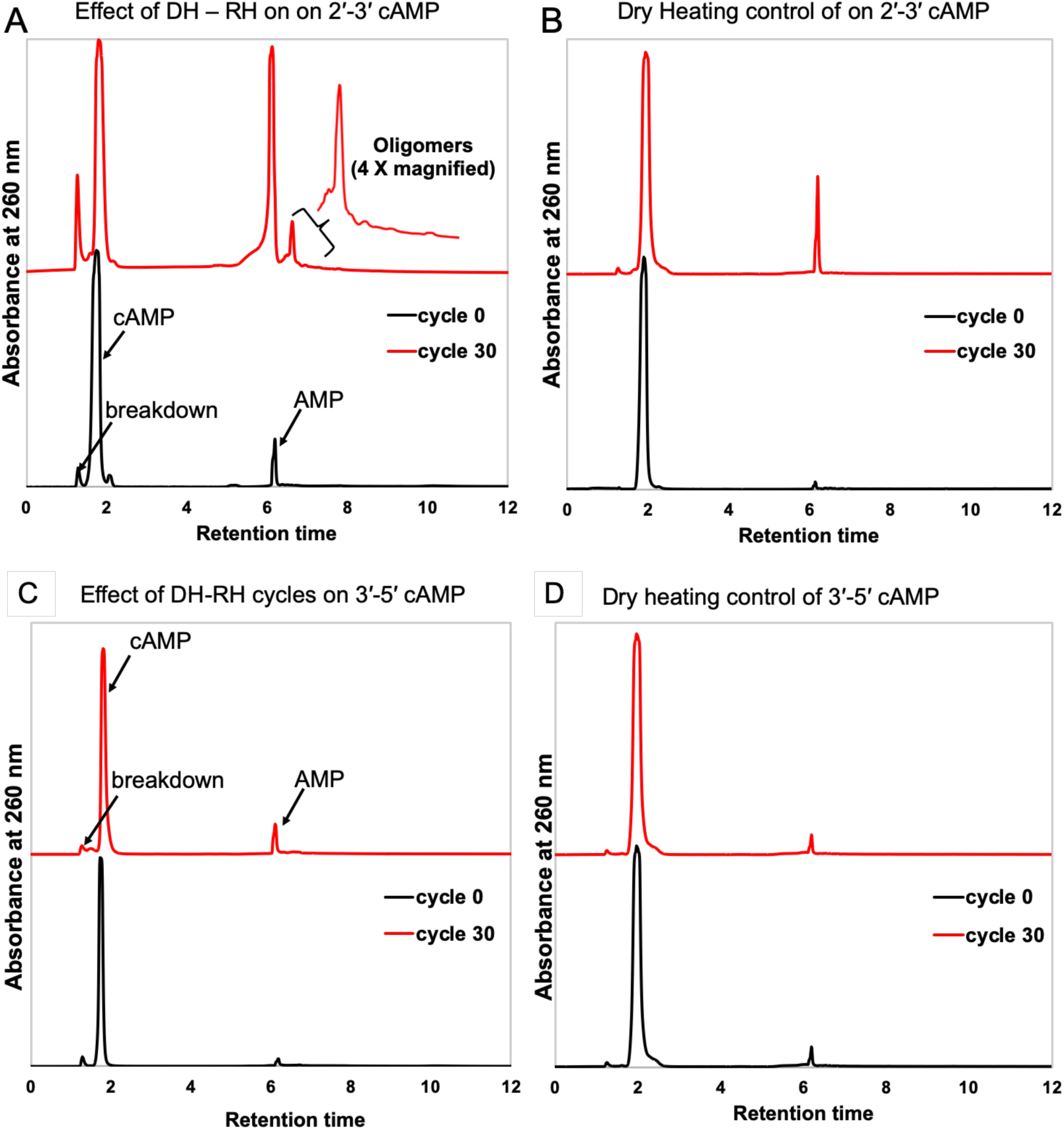
Effect of DH-RH cycles on the oligomerization of cAMP. Representative HPLC chromatograms for oligomerization reactions of 2′, 3′ cAMP and 3′, 5′ cAMP subjected to thirty DH-RH cycles (Panel A and C, respectively); Oligomerization of 2′, 3′ cAMP and 3′, 5′ cAMP, but only under prolonged dry heating (Panel B and D, respectively). The black trace depicts the chromatogram for cycle 0, which was collected at time zero, and the red trace shows the chromatogram for reaction time point equivalent to thirty days. Y-axis shows the absorbance at 260 nm while the X-axis shows time in minutes.

Time of Flight–Mass Spectrometry (TOF-MS) was performed in order to further confirm and characterize the various species that were observed in the HPLC chromatograms (Method 4.2.6). Fragmentation was induced for the precursor molecule using the TOF-MS/MS mode. Figure 2 shows representative TOF-MS and TOF-MS/MS spectrums that were observed in Electro Spray Ionization (ESI) positive mode for H^+^ adducts of AMP linear dimer (Panels A and B), along with the corresponding structure (Panel C), AMP linear trimer (Panels D and E), and the corresponding structure (Panel F), and AMP linear tetramer (Panels G and H), and the corresponding structure (Panel I). All the observed masses have been summarized in Table 1, with the errors being well within the acceptable range (give or take +/- 5 parts per million (ppm)). The fragmentation of AMP linear dimer with a monoisotopic mass of 677.123 Da, upon fragmentation yielded an intense fragment at 348.067 Da, and another fragment at 542.064 Da (Figure 2, Panel A and B). AMP linear trimer with a monoisotopic mass of 1006.170 fragmented into two intense fragments of 348.067 Da, and 677.112 Da (Figure 2, Panel D and E). AMP linear tetramer with a monoisotopic mass of 1335.238, fragmented into two intense fragments, of 348.074 Da, and 677.12 Da (Figure 2, Panel G and H). The fragmentation fingerprint of the parent molecule, its ppm accuracy, and the elution time were used in conjunction to confirm the presence of the target molecules. The TOF-MS spectrum for the 2′, 3′ cAMP reaction shows the presence of intact linear dimer within four DH-RH cycles, and intact linear trimer and tetramer after ten DH-RH cycles (Figure 6). Nevertheless, 3′, 5′ cAMP yielded oligomer upto linear intact dimer even after twenty DH-RH cycles. The reduced oligomerization potential of 3′, 5′ cAMP could possibly stem from its higher intrinsic stability due to the presence of reduced strain on its six-membered ring as compared to the five-membered ring of 2′, 3′ cAMP where the ring strain is more pronounced (Khorana et al. 1957).

**Table 1:** Summary of the masses observed in the nonenzymatic oligomerization reactions of cAMP and cCMP reactions. * depicts that the species was observed in TOF-MS but not seen in TOF-MS-MS analysis.

|  | Species | Expected mass | Observed mass | Error (ppm) |  | Species | Expected mass | Observed mass | Error (ppm) |
| --- | --- | --- | --- | --- | --- | --- | --- | --- | --- |
| 2'-3' cNMP | cAMP | 330.05 | 330.05 | -0.90 |  | cCMP | 306.04 | 306.04 | -1.30 |
|  | AMP dimer | 677.12 | 677.12 | 2.51 |  | CMP dimer | 629.10 | 629.10 | -0.15 |
|  | AMP trimer | 1006.17 | 1006.17 | 1.78 |  | CMP trimer | 934.14 | 934.14 | 1.92 |
|  | AMP tetramer | 1335.22 | 1335.23 | 2.62 |  | CMP tetramer * | 1239.18 | 1239.18 | 4.59 |
| 3'-5' cNMP | cAMP | 330.05 | 330.05 | -1.51 |  | cCMP | 306.04 | 306.04 | -0.98 |
|  | AMP dimer | 677.12 | 677.12 | 1.77 |  | CMP dimer | 629.10 | 629.10 | 1.74 |

**Figure 2:**
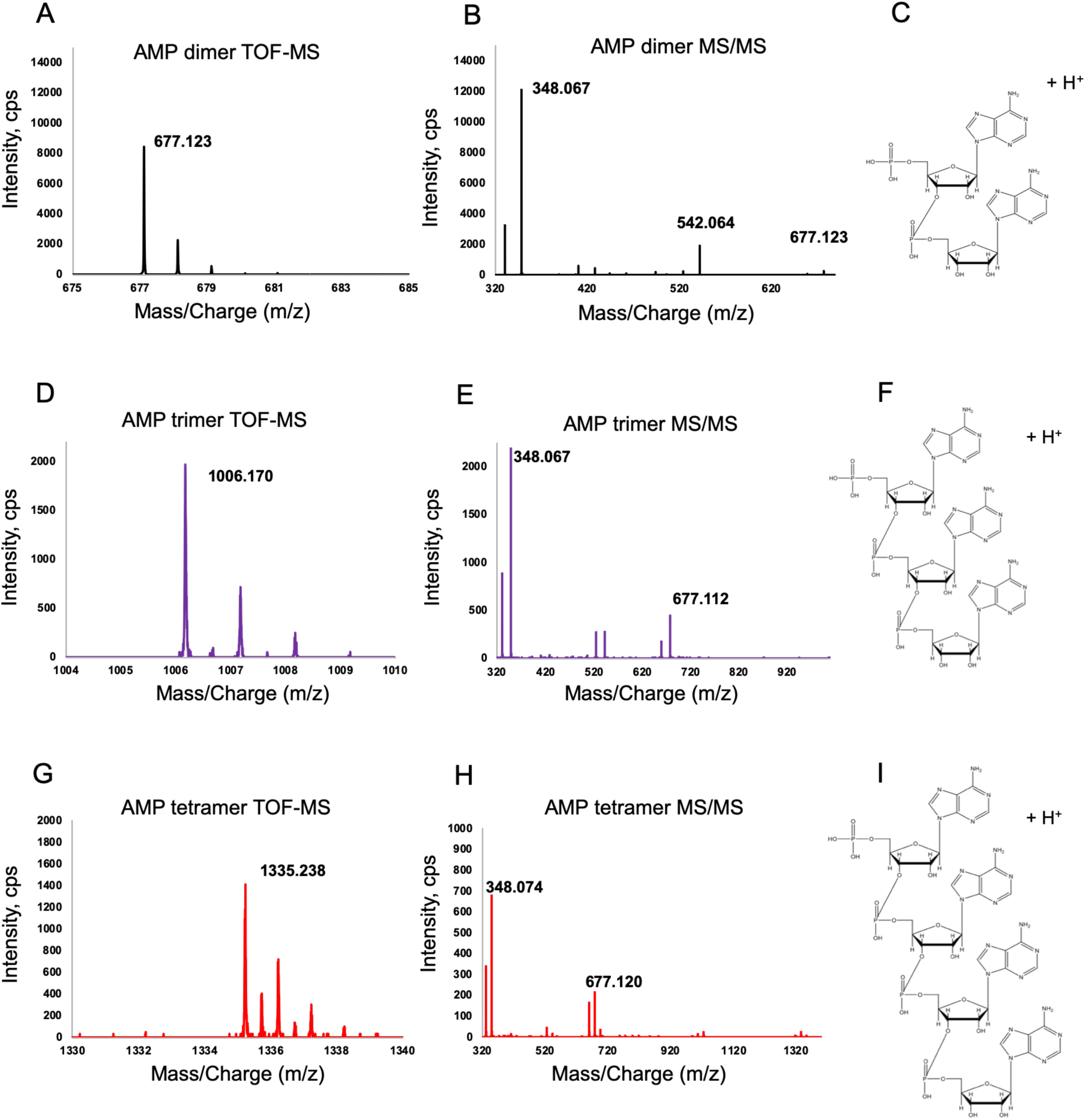
Representative TOF-MS and TOF-MS/MS spectra in ESI positive mode for cAMP reactions. From left to right: the TOF-MS spectrum, TOF-MS/MS spectrum, and the representative structure of different AMP oligomer species observed. The species correspond to H^+^ adducts of AMP linear dimer (A, B, and C) with a monoisotopic mass of 677.12 Da, AMP linear trimer (D, E, and F) with a monoisotopic mass of 1006.17 Da, and AMP linear tetramer (G, H, and I) with a monoisotopic mass of 1335.23 Da, respectively.

**Figure 3:**
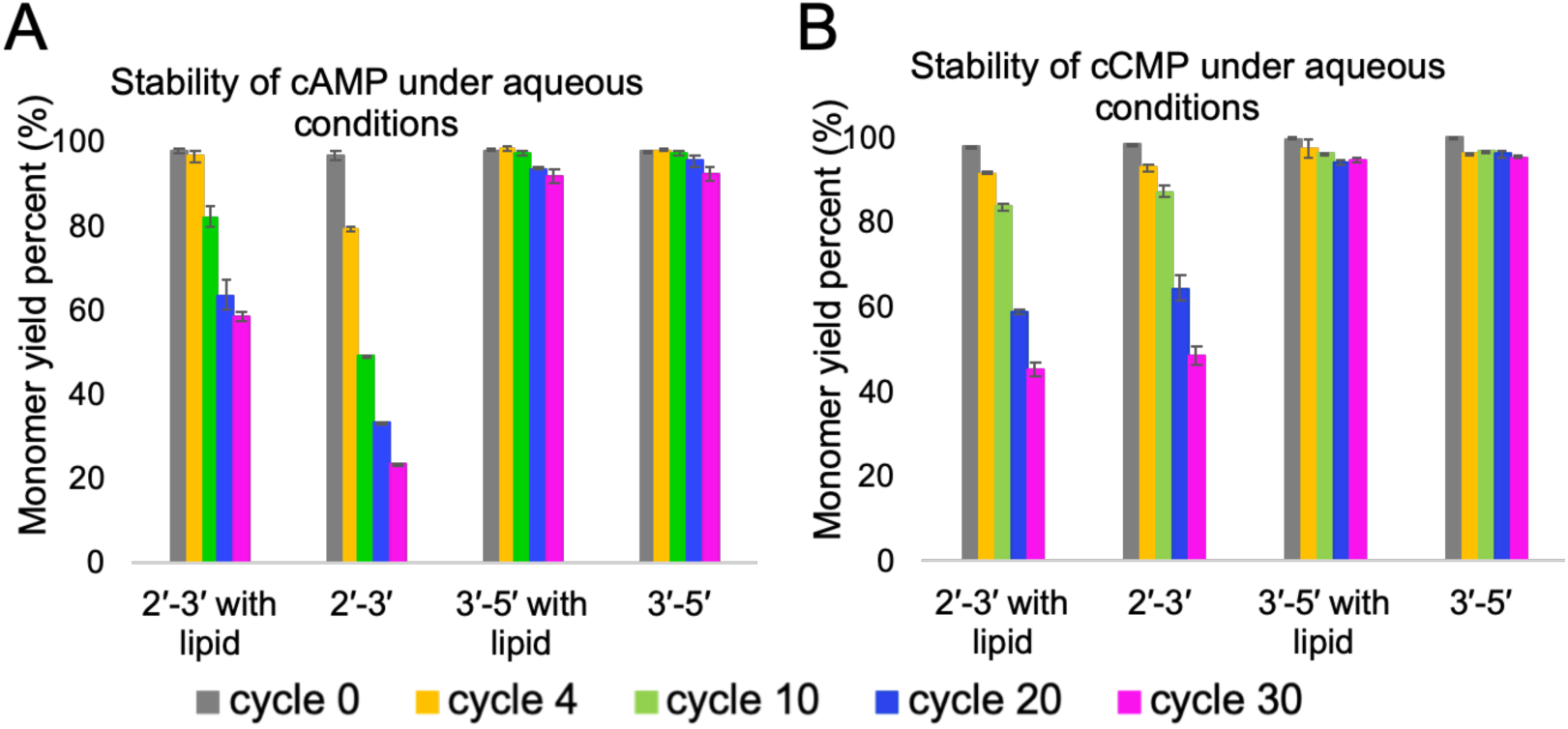
The stability of cyclic nucleotides in enzyme-free oligomerization reactions, in both, without lipid and lipid-assisted reactions. The graphs show the stability of the cAMP (Panel A) and cCMP (Panel B), during the reactions carried out under DH-RH regimen, either in the presence or absence of lipids. X-axis shows different reactions with indicated cNMP. Different colors depict different DH-RH cycles. Y-axis shows monomer yield percent normalized to that present in cycle zero. Error bars indicate s.d.

**Figure 4:**
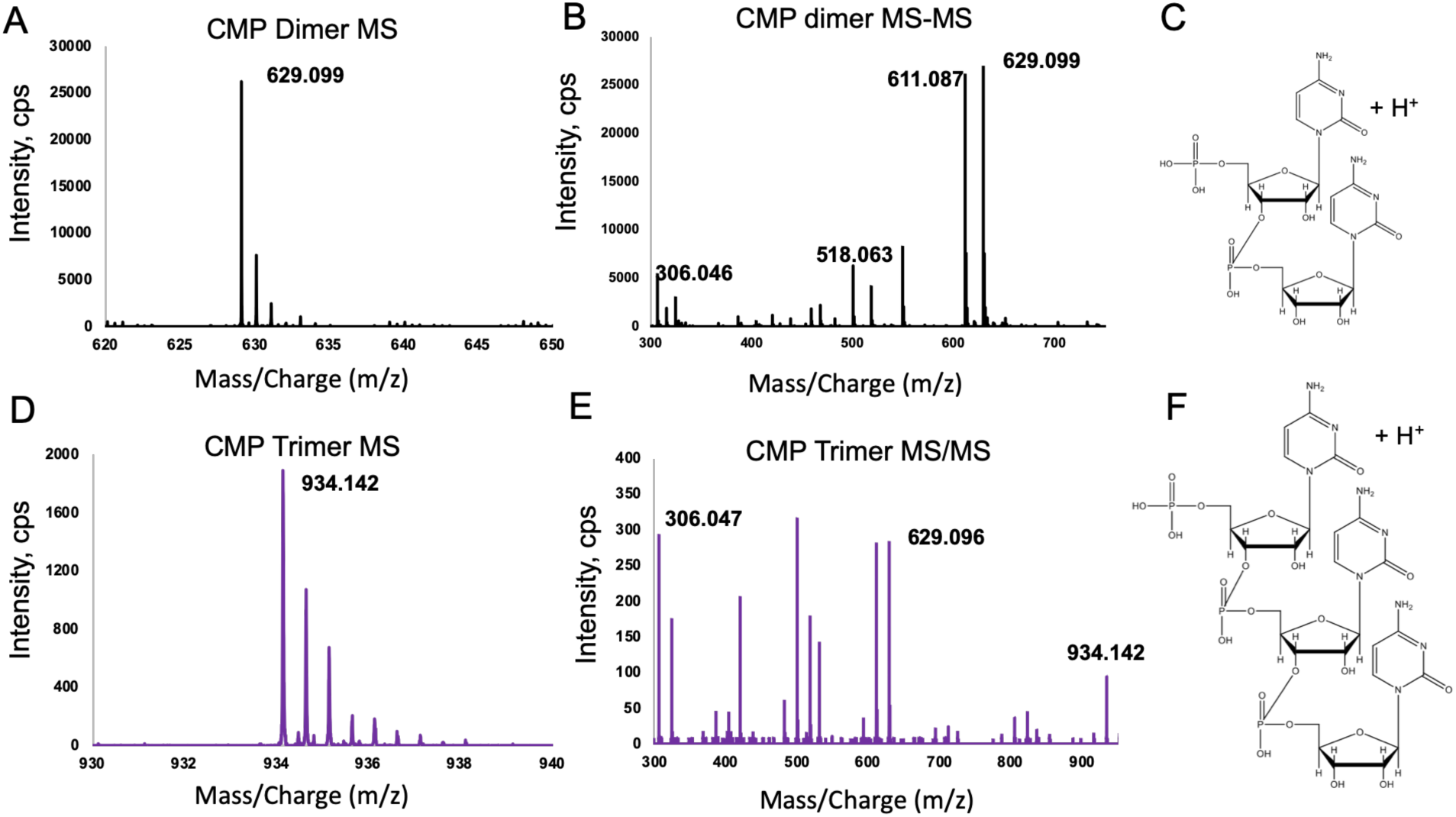
Representative spectra of TOF-MS and TOF-MS/MS in ESI positive mode for cCMP based reactions. The figure represents the TOF-MS spectrum, TOF-MS/MS spectrum, and the structures that were observed for the H^+^ adducts of CMP linear dimer (A, B, and C) with monoisotopic mass 629.1 Da, and CMP linear trimer (D, E, and F) with monoisotopic mass 934.14 Da, respectively.

**Figure 5:**
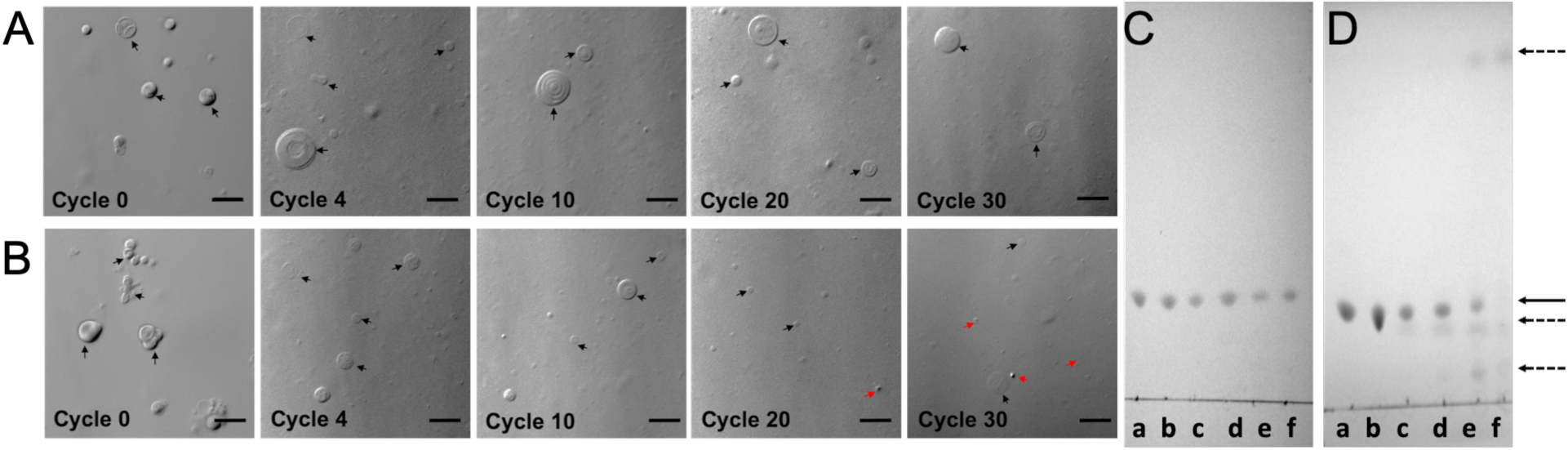
The stability of POPC vesicles under our reaction conditions. The stability of POPC vesicles under oligomerization reaction conditions with increasing DH-RH cycles (left to right), viz. cycle 0, cycle 4, cycle 10, cycle 20, cycle 30, either under aqueous conditions (Panel A) or in Panamic water sample (Panel B). The black and red arrows indicate vesicles and aggregates (collapsed vesicles), respectively. The scale bar in all the images is 10 microns. The chemical stability of POPC molecules under DH-RH conditions in nanopure water (Panel C) and Panamic water (panel D). Different lanes from left to right i.e. a to f, indicate POPC control, the reactions with corresponding to DH-RH cycle 0, cycle 4, cycle 10, cycle 20, and cycle 30, respectively. Solid black arrow indicates the band corresponding to POPC, dashed black arrows indicate new bands that appear corresponding to degradation products.

**Figure 6:**
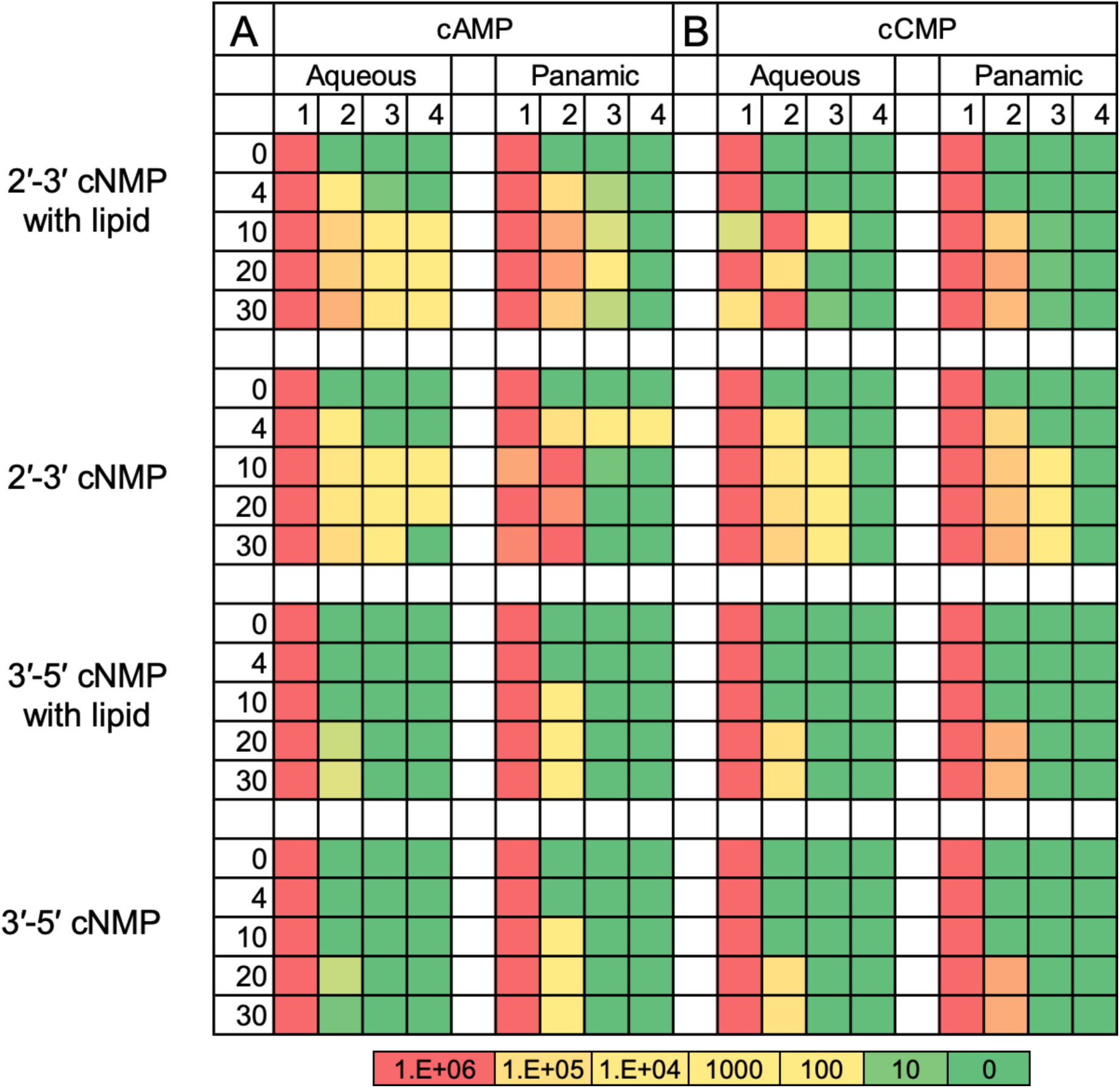
Heat map of all the species observed in enzyme-free oligomerization reactions using LC-MS/MS. Panel A and B shows the species observed in different reactions of cAMP, and cCMP respectively as mentioned towards the left of the figure. The numbers viz. 0, 4, 10, 20 and 30, on the left axis of the heat map indicates the number of DH-RH cycles. Lanes 1 to 4 shows the presence of cNMP, dimer, trimer, and tetramer, respectively. Each block depicts a single reaction with rows and columns, signifying different time points and different species observed, respectively.

### 2.2. Effect of lipids on the stability of cAMP isomers under DH-RH conditions

Lipids have been demonstrated to encapsulate macromolecules under DH-RH cycles (Deamer and Barchfeld 1982), and are also known to interact better with certain molecules that might be present in the bulk solution (Black et al. 2013b; Sasidharan et al. 2019), consequently protecting them from the harsh environmental conditions of an early Earth. In the context of RNA, lipids have been shown to also facilitate the nonenzymatic oligomerization of NMPs to form RNA-like polymers (abasic oligomers) (Rajamani et al. 2008; Mungi et al. 2015). Given this, we set out to elucidate the plausible effect of lipids on the enzyme-free oligomerization of cAMP. The reactions were performed in the presence of 1-palmitoyl, 2-oleoyl phosphatidylcholine (POPC), a phospholipid that has been used a proxy for lipid in several previous studies. 1 mM of POPC was added to each reaction containing 5 mM corresponding cAMP to maintain a constant ratio of 5:1 of nucleotide is to lipid. The reaction conditions and subsequent procedures were kept similar to those mentioned in the previous section (90 °C, thirty DH-RH cycles with 24 hours per cycle) (Methods 4.2.3). In order to get an estimate of the amount of starting monomer in the HPLC runs, the area under, both, linear AMP and cAMP peaks were taken into consideration, as cAMP can readily convert to AMP under aqueous conditions (Suárez-Marina et al. 2019). Figure 3 (Panel A) shows the semi-quantitiave HPLC results obtained for the stability of the cAMP starting monomer in the reactions that were carried out under DH-RH regimen in the absence of lipids. The amount of cAMP decreased with increasing DH-RH cycles. In the 2′, 3′ cAMP reactions, cAMP reduces to 80% of the starting concentration in about four DH-RH cycles, further decreasing to 50 %, 30 % and 20% in ten, twenty and thirty cycles, respectively. However, in lipid-assisted reactions, the starting monomer was more stable, upto 60 % persisted even after thirty DH-RH cycles. In case of 3′, 5′ cAMP, only 5 % of the starting monomer underwent breakdown even after thirty DH-RH cycles. The results suggested that the starting monomer seemed much more stable in the lipid-assisted reactions, particularly the one involving 2′, 3′ cAMP. However, the protecting effect of lipid was not evident in the reactions with 3′, 5′ cAMP.

### 2.3. Nonenzymatic oligomerization reactions using cCMP

As mentioned earlier, purines are known to stack better in comparison to pyrimidines and thus can oligomerize faster (Giovanna et al. 2009). In order to investigate how the oligomerization of cyclic nucleotides differs for a purine as compared to a pyrimidine nucleobase, reactions were carried out using cyclic cytosine monophosphate (cCMP). The reactant concentrations and reaction conditions including DH-RH cycling (90 °C, thirty DH-RH cycles with 24 hours per cycle), were kept similar to what was used in the cAMP reactions. Both of the cyclic forms i.e. 2′, 3′-cCMP and 3′, 5′-cCMP were investigated for their nonenzymatic oligomerization potential. The effect of lipid (POPC) was also investigated in these reactions. The resultant products were analyzed using HPLC.

Figure 3 (Panel B) shows the graph of the semi-quantitation done on the HPLC data, which corresponds to the stability of starting monomer (accounting for both cCMP and CMP). The results suggested that the amount of the starting monomer decreased with increasing DH-RH cycles. The quantity of 2’, 3’ cCMP reduced by 10 %, 20 %, 40 % and 55 % in four, ten, twenty and thirty days of DH-RH cycling, respectively, in both the without lipid and lipid-assisted reactions. 3′, 5′ cCMP was found to be more stable than the 2’, 3’ cCMP under our reaction conditions, similar to what was observed with cAMP isomers. In all, the concentration reduced to 95 % of its initial concentration in thirty DH-RH cycles. The lipid did not seem to have any influence on the cCMP reactions, both, in the case of 2′ -3′ cCMP and 3′ -5′ cCMP reactions. As alluded to earlier, the protecting effect of lipid on a particular molecule could stem from either its entrapment inside a vesicle/within bilayers, or by directly interacting with the amphiphilic surfaces. Few studies have reported that purines have a much better tendency to interact with a vesicle than pyrimidines (Black et al. 2013a; Sasidharan et al. 2019). The reduced interaction of pyrimidine nucleobases/nucleotides with lipid can possibly explain why there probably is no noticeable effect of the lipid in the case of nonenzymatic oligomerization reactions involving cCMP.

Mass analysis of the cCMP oligomerization reaction products was performed. The parameters for TOF-MS and TOF-MS/MS were kept similar to that of the cAMP based reactions (Method 4.2.6). Figure 4 shows a representative TOF-MS spectra and TOF-MS/MS spectra in ESI positive mode that were observed for the H^+^ adduct of CMP linear dimer (Panels A and B), along with the corresponding structure (Panel C), and H^+^ adduct of CMP linear trimer (Panels D and E) and the corresponding structure (Panel F). As seen in Figure 4, the CMP linear dimer with a monoisotopic mass of 629.099 Da, fragmented into daughter fragments with masses 611.087 Da, 306.046 Da, and 518.063 Da. CMP linear trimer with a monoisotopic mass of 934.142, fragmented into 306.047 Da, and 629.096 Da, respectively. All the observed masses with their respective ppm errors are summarized in Table 1. The TOF-MS and MS/MS analysis for the 2′,3′ cCMP and 3′, 5′ cCMP reactions show the presence of linear trimer and linear dimer, respectively, within ten DH-RH cycles (Figure 6). To summarize, cCMP could oligomerize under these DH-RH cycling conditions, but it, however, was to a lesser extent than cAMP.

### 2.4. Oligomerization reactions of cyclic nucleotides under analogue conditions

To obtain a more realistic picture of the oligomerization phenomenon involving cyclic nucleotides, the enzyme-free oligomerization reactions were also performed using water samples collected from a hot spring. Few previous studies have demonstrated that the presence of certain metal ions can catalyze nonenzymatic oligomerization reactions of nucleotides (Lailach et al. 1968; Ferris 2005; Joshi et al. 2012). Thus, metal ion co-solutes that are present in a hot spring could also potentially affect the nonenzymatic oligomerization of nucleotides. Nonetheless, high amounts of salt is known to be detrimental for RNA (Blasko and Bruice 1999). Panamic hot spring water from Joshi et al. study was chosen, to study the afore-mentioned oligomerization reactions (Joshi et al. 2017) in detail. This site was studied as part of Spaceward Bound expedition to Ladakh, an astrobiologically relevant site in the northern part of India (Pandey et al. 2019). It is considered as early Earth and Martian analogue site due to its topological feature. The reaction conditions and other parameters were kept similar to what was described for the aforesaid reactions that were studied under aqueous conditions i.e. thirty DH-RH cycles, each cycle consists of 24 hours at 90 °C. The native pH of the hot spring water sample was ~8 - 8.5, which was kept as is in the experiments undertaken.

Figure 6 gives a comparative summary of the different oligomeric species that were observed both in the presence and absence of lipid in aqueous as well as analogue reactions, when using cAMP and cCMP as starting monomers. All the oligomers were characterized using LC-MS and MS/MS analysis. 2′, 3′ cAMP based reactions yielded oligomers up to intact AMP linear tetramer after four DH-RH cycles. However, it seemed to degrade after ten cycles of DH-RH. AMP linear trimer was also observed to disappear after twenty DH-RH cycles. On the contrary, in lipid-assisted reactions, 2′, 3′ cAMP oligomerized to yield linear trimer in four days of DH-RH cycling and persisted even after thirty DH-RH cycles, indicating the stabilizing effect of lipid. However, in the case of 3′, 5′ cAMP reaction, only a dimer was observed after ten DH-RH cycles, both, in the presence and absence of lipid. Therefore, the presence of lipid did not seem to make any significant difference in this case. In 2′, 3′ cCMP lipid-assisted and without lipid reactions, CMP linear trimer was observed after ten DH-RH cycles, which was stable throughout the reaction i.e. till thirty cycles. However, the relative amount of trimer was less in lipid-assisted reaction. The experiments with 3′, 5′ cCMP yielded linear dimer in twenty DH-RH cycles. In general, the metal ions and other co-solutes present in hot spring water (like silicates etc.) seemed to increase the rate of oligomerization reactions, albeit destabilizing the resultant oligomers too.

### 2.5. Stability of POPC vesicles under multiple DH-RH cycles in aqueous and analogue conditions

Our results demonstrated that the presence of POPC could indeed enhance the stability of the starting monomers under DH-RH conditions, which corroborates with previous studies (Rajamani et al. 2008; Mungi et al. 2015). As discussed by Damer and Deamer, lipid bilayers can do so by trapping the substrate molecules in the interlamellar spaces during the dry phase. Coincidentally, this also provides a concentration effect thereby facilitating forward reactions like oligomerization. Furthermore, the lipids would also encapsulate the resultant oligomers upon rehydration (Damer and Deamer 2015), thus protecting them from back reactions like hydrolysis, which leads to their accumulation over multiple cycles. However, to the best of our knowledge, none of the previous studies have examined the stability of the POPC vesicles under multiple DH-RH cycles for a prolonged duration, like thirty days, as was done in this study. In order to evaluate whether the lipid vesicles are stable under our reaction conditions, 1 mM of POPC was used and DH-RH cycling was performed, both, under aqueous conditions and in the Panamic water sample, at pH 8 for thirty days. The stability of the POPC vesicles was monitored by observing them under microscope after different number of DH-RH cycles, using Differential interference contrast (DIC) microscopy (Method 4.2.2 and 4.2.4). Intact vesicles were observed under aqueous conditions even after thirty DH-RH cycles (Figure 5, Panel A). On the contrary, small aggregates (collapsed vesicles) were observed along with vesicles after twenty and thirty DH-RH cycles in the Panamic water sample (Figure 5, Panel B). Nonetheless, an important aspect to consider here is that mere presence of vesicles does not necessarily have to be a direct reflection of the molecular stability of POPC. Previous literature in the field has demonstrated that the fatty acid chains can also self-assemble into vesicles around the pH that is near or equal to their pKa (Monnard et al. 2015). Oleic acid has a pKa around 8-8.5. Consequently, at our reaction pH i.e. 8, if POPC is degrading to its components, the oleic acid can potentially form vesicles. Thus, the vesicles observed could also be a false positive indication for the stability of POPC. Therefore, the chemical stability of POPC molecules was also evaluated using Thin Layer Chromatography (TLC), in experiments involving the aqueous reaction samples and the Panamic water samples (Methods 4.2.7). The stability of POPC decreased with increasing number of DH-RH cycles (Figure 5; panels C and D), under both the reactions conditions. It seemed to be comparatively more stable under aqueous conditions (Figure 5, Panel C) than in the Panamic water sample reaction (Figure 5, Panel D). This was evident from the intensity of TLC band corresponding to POPC (band corresponding to black solid arrow), which disappeared by thirty DH-RH cycles in Panamic water. With increasing DH-RH cycles, there were also some new bands that were observed, which correspond to the degradation products. These were mainly fatty acids (palmitic acid and oleic acid) and monoacylated phospholipid, which was confirmed by High Resolution Mass-Spectrometry (HRMS). As mentioned earlier, fatty acids (oleic acid) on their own have a tendency to form vesicles at pH 8. So, the vesicles observed in the Panamic water sample after 30 cycles of DH-RH microscopy (Figure 5, Panel B) could also be pure fatty acid vesicles or a mixed mebrane composed of POPC and fatty acid.

## 3. Discussion

Cyclic nucleotides have been hypothesized to undergo oligomerization via a base-catalyzed reaction (Costanzo et al. 2012b). Our study demonstrates that prebiotically relevant DH-RH cycles can indeed facilitate the enzyme-free oligomerization of, both, 2′, 3′ and 3′, 5′ cyclic nucleotides for a purine and a pyrimidine based system. Subsequent cycles of DH-RH can act as a kinetic trap by pushing the reaction towards oligomerization, thereby facilitating formation of longer oligomers over multiple cycles. Although, the rehydration phase is necessary for effective collisions to occur between monomers and /or the growing oligomers, it can also in principle, enhance the back hydrolysis of oligomers that are already present. Therefore, there is always a dynamical interplay of oligomerization and hydrolysis in these geological settings.

Under DH-RH conditions, 3′, 5′ cNMP was observed to be more stable as compared to 2′, 3′ cNMP for both, purine (cAMP) and pyrimidine (cCMP) based systems. Presence of lipids also seemed to increase the stability of the starting monomers. This was especially pronounced in the case of 2′, 3′ cAMP, indicating a protecting effect coming from the lipid. However, there was no prominent effect observed in the case of 3′, 5′ cAMP. Oligomers till tetramers were observed by mass analysis, in the case of, both, 2′, 3′ cAMP and 2′, 3′ cCMP oligomerization reactions (Table 1). On the contrary, when 3′, 5′ cNMPs were used as the starting monomer, oligomers only up to dimers were observed. This could be attributed to the high intrinsic stability of 3′, 5′ cNMPs. 3′, 5′ cNMP possess a six-membered cyclic ring, which is thermodynamically more stable than the five-membered cyclic ring that is present in 2′, 3′ cNMPs (Khorana et al. 1957). The role of different metal ions, and how heterogeneity of the prebiotic pool can affect the efficiency of oligomerization, was also investigated. Towards this, reactions were carried out in a hot spring water sample obtained from Panamic hot spring in Ladakh. In these analogue sample-based reactions, the oligomers seemed to form more readily, as was observed for e.g. in the reactions corresponding to 2′, 3′ cAMP (tetramer was observed after four days of DH-RH cycles) (Table 2, Figure 6; Panel A). However, the oligomers formed were found to be more susceptible to hydrolysis too, potentially because of the presence of different ions. This is in contrast to what was seen in lipid-aided analogue reactions wherein although oligomers up to trimers were formed in these reactions, the resultant oligomers were stable even after thirty days of DH-RH cycling. Interestingly, 2′, 3′ cNMP upon oligomerization is thought to potentially yield 2′, 5′ as well as 3′, 5′ linked phosphodiester bonds (Costanzo et al. 2016a). On the contrary, 3′, 5′ cNMP can oligomerize to yield only 3′, 5′ linked phosphodiester bonds. Previous literature have suggested that 2′, 5′ phosphodiester linkages are more prone to hydrolysis as compared to 3′, 5′ linkages (Usher and McHale 1976). Our study emphasizes the fact that, though the speed of oligomerization of 3′, 5′ cNMPs is compromised, the synthesized oligomers are found to be relatively more stable than that of 2′, 3′ cNMPs (Figure 3 and 6). This reduced stability of products from 2′, 3′ cNMPs oligomerization could be due to the formation of 2′, 5′ linked oligomers, which are more prone to hydrolyze in comparison to the 3′, 5′ linked oligomers. The higher stability of oligomers corresponding to 3′, 5′cNMP reactions, in both, nanopure water as well as in hot spring water (a proxy for prebiotically pertinent conditions), could be indicative for how selection of 3′, 5′ phosphodiester linkages might have been driven by such relevant selection pressures. In case of cCMP reactions, similar extent of oligomerization was observed in reactions that were carried out under aqueous conditions and Panamic water (Figure 6; Panel B). The lower potential of cCMP to oligomerize, both, under aqueous conditions and in Panamic water, could potentially be due to the higher stability of cCMP than cAMP (Gerlt et al. 1974). The higher stability of cCMP reduces its tendency to oligomerize as it requires ring opening up of cyclic phosphodiester linkage. Our results seemed to indicate that the presence of ions in the analogue water sample, could effectively act as a selection pressure playing an important role in shaping the evolutionary landscape of RNA World.

On investigating the stability of lipid under our reaction conditions, microscopic analysis suggested that lipid vesicles are stable up to thirty DH-RH cycles, both, under aqueous conditions and in Panamic water. Additionally, even though POPC molecules seemed to be stable under aqueous conditions (nanopure water) during the course of our reaction time period, it seemed to degrade in the Panamic water sample (Figure 5, Panel D).

On the whole, this study showed the formation of intact informational oligomers under DH-RH conditions using cyclic nucleotides. To our knowledge, very few studies have tried to delineate the oligomerization potential of 2′, 3′ cyclic isomer (Tapiero and Nagyvary 1971; Verlander et al. 1973) or a pyrimidine nucleotide (Tapiero and Nagyvary 1971; Costanzo et al. 2017). The present study illustrates the enzyme-free oligomerization resulting in the formation of intact oligomers in both a purine- (cAMP) and a pyrimidine-based (cCMP) system, under laboratory-simulated as well as analogue conditions. The stability of all the systems was systematically demonstrated. In addition, the protecting effect of lipid was also shown on the aforementioned reactions. It has been reported that short oligomers (up to trimers) can enhance the rate of template-directed replication by three folds (Kaiser and Richert 2013; Li et al. 2017; Tam et al. 2017; O’Flaherty et al. 2018). Thus, the intact oligomers reported in the present study, can, potentially assist the propagation of information in template-directed replication scenarios. Altogether, this study provides a comprehensive and robust means by which enzyme-free oligomerization of intrinsically activated, prebiotically pertinent nucleotides might have occurred and evolved in a heterogenous prebiotic soup.

## 4. Experimental section

### 4.1. Materials

The monosodium salts of all four cyclic monophosphates viz. adenosine 3′, 5′ -cyclic monophosphate (3′, 5′ -cAMP), adenosine 2′, 3′ -cyclic monophosphate (2′, 3′ -cAMP), cytosine 3′, 5′-cyclic monophosphate (3′, 5′-cCMP), cytosine 2′, 3′-cyclic monophosphate (2′, 3′-cCMP), adenosine monophosphate (AMP) linear dimer, trimer, and tetramer were purchased from Sigma-Aldrich (Bangalore, India) and used without further purification. 1-palmitoyl-2-oleoyl-sn-glycero-3-phosphocholine (POPC) was purchased from Avanti Polar Lipids Inc. (Alabaster, AL, USA). All other reagents used were of analytical grade and purchased from Sigma-Aldrich (Bangalore, India). TLC Silica gel 60 F_254_ was purchased from Merck (EMD Millipore corporation, Billerica, USA).

### 4.2. Methods

#### 4.2.1. Simulating early-Earth like conditions

Early Earth-like conditions were simulated using a bench-top heating block that was maintained at high temperatures i.e. 90 °C as described by Mungi et.al. in 2015. The oligomerization reactions were carried out in 20 ml glass vials with their caps fitted with PTFE septa purchased from Chemglass (Vineland, NJ, USA). Anaerobic environment was maintained by delivering gentle flow of carbon dioxide into the vials through two PEEK tubings of about 1–1.5 inches, one acting as an inlet and another as an outlet for the gas.

#### 4.2.2. Vesicle solution preparation

The vesicle solutions were prepared by drying the required amount of chloroform solution (with strength 25mg/ml) of POPC under nitrogen gas flow to prepare a dry lipid film. It was then kept under vacuum for five to six hours to make sure that no trace amount of chloroform remained. Subsequently, nanopure water was used to rehydrate the thin film to form the vesicles.

#### 4.2.3. Nonenzymatic oligomerization reaction conditions and procedure

A typical reaction mixture consisted of 5 mM cyclic nucleotide in nanopure water with pH ~8.5 or hot spring water sample. In lipid-assisted reactions one millimolar of POPC was added to each reaction to maintain a constant ratio of 5:1 for nucleotide is to lipid. The lipid concentration was chosen based on a previous study in our lab (Mungi et al. 2015). The samples were rehydrated with nanopure water, and mixed properly followed by a prolonged dehydration phase during each DH-RH cycle, with 24 hours per cycle. The rehydrated samples were collected at different periods. The lipid was extracted from the rehydrated sample using pre-standardized butanol-hexane extraction procedure. The aqueous portion containing nucleotides was collected, and analyzed using HPLC and LC-MS/MS.

#### 4.2.4. Microscopic analysis

10 μL of the reaction sample containing only one mM of POPC, was placed on a glass slide which was then evenly spread and covered with a 18×18 mm coverslip. This was then air sealed by covering its four sides with liquid paraffin. This was done in order to decrease the motion of solution. The slides prepared were then observed under 40X magnification using a Differential Interference Contrast (DIC) microscope AxioImager Z1 (Carl Zeiss, Germany), (NA = 0.75) to check for the presence of lipid vesicles.

#### 4.2.5. HPLC analysis

HPLC analysis was performed using Infinity series 1260 HPLC instrument (Agilent Technologies, Santa Clara, CA, USA). The reaction products, after removing the lipid molecules using previously standardized Butanol-Hexane extraction (Mungi et al. 2015), were loaded onto HPLC. The molecules were separated by an anion-exchange column viz. DNAPac PA 200 (Thermo Scientific, Sunnyvale, CA, USA). It separates the molecules based on their interaction with the column through phosphate moiety, thus providing single-nucleotide resolution. The separation method was standardized using a gradient of sodium perchlorate as eluting solvent and commercially available cAMP monomer, linear AMP monomer, dimer, trimer and tetramer as standards. A photo Diode-Array Detector (DAD) was used to detect analytes at 260nm, using a highly sensitive 60 mm flow cell. The separation between a dead volume peak (which was later characterized as open ring structures lacking phosphate moiety i.e. adenosine or adenine by mass), cAMP monomer, linear AMP monomer, and oligomers was observed in a typical HPLC chromatogram. The analytes were semi-quantified by measuring the area under corresponding peaks separated through column. Although the semi-quantified area can be due to the presence of various species with the same charge, it can still provide us with the distribution of species in the reaction after a particular DH-RH cycle.

#### 4.2.6. Mass analysis

Mass analysis of the samples was performed on Sciex X500R QTOF mass spectrometer (MS) fitted with an Exion-LC series UHPLC (Sciex, CA, USA) using Information Dependent Acquisition (IDA) scanning method. The acquired data was analyzed using the Sciex OS software (Sciex, CA, USA; University of Florida, FL, USA). The crude reaction mixture was separated on Zorbax C8 column (dimensions: 4.6 × 150 mm, 3.6 μm particle size) (Thermo Scientific, Sunnyvale, CA, USA) fitted with its guard column. A gradient of MilliQ containing 0.1% formic acid and acetonitrile containing 0.1% formic acid was used to separate the oligomers. All the mass acquisition was performed using Electron spray ionization (ESI) with the following parameters: turbo spray ion source, medium collision gas, curtain gas = 30 L/min, ion spray voltage = 5500 V (positive mode), at 500 °C. TOF-MS acquisition was done at declustering potential as 80 V with 20 V as spread and using 10 V collision energy. To perform TOF MS-MS analysis, 50 V collision energy with 20 V spread was used. As the mass acquisition was carried out in positive mode, the observed masses correspond to the mass of the H^+^-adduct of the parent molecule. The presence of a specific species/molecule was confirmed by the presence of precursor mass within 5 ppm error range as well as its fragmentation pattern.

#### 4.2.7. TLC (Thin Layer Chromatography) of POPC

TLC analysis for evaluating the chemical stability of POPC molecules was performed using TLC Silica gel 60 F_254_ (Merck, EMD Millipore corporation, Billerica, USA). A mixture of chloroform, methanol and water in 65:25:4 was used as the mobile phase. The TLC chamber (Latch-lid TLC developing chambers for 10 × 10 cm plates, Sigma-Aldrich, Bangalore, India) was lined with filter paper and equilibrated with the solvent system before use. The TLC plates (dimension 5 × 10 cm) were pre-run with the mobile phase in order to eliminate interference coming from any intrinsic contaminant. It was then dried, and 6 µl of each sample withdrawn at different DH-RH cycles (cycle 0, cycle 4, cycle 10, cycle 20 and cycle 30 in lanes b, c, d, e, and f, respectively) along with POPC as a control (lane a), was spotted 0.8 cm apart to each other and 2 cm above the bottom edge of TLC plate. The dried TLC plates were then chromatographed in the equilibrated chamber. Following the chromatography, the TLC plates were dried and kept in the pre-saturated tank containing iodine vapors. The plates were then immediately scanned on a Syngene G-Box Chemi-XRQ gel documentation system (Syngene, Cambridge, UK).

## Acknowledgments

The authors wish to acknowledge the Mass spectrometry facility (co-funded by DST-FIST and IISER Pune) and the Microscopy facility at IISER Pune. SR acknowledges Department of Biotechnology, Govt. of India [BT/PR19201/BRB/10/1532/2016] for extramural funding. SD acknowledges the research fellowship received from CSIR, Govt. of India. SS acknowledges the research fellowship received from IISER Pune. We are grateful to Dr. Niraja Bapat and Mr. Manesh P. Joshi for their critical inputs on the manuscript.

